# Primitive purine biosynthesis connects ancient geochemistry to modern metabolism

**DOI:** 10.1101/2022.10.07.511356

**Authors:** Joshua E. Goldford, Harrison B. Smith, Liam M. Longo, Boswell A. Wing, Shawn E. McGlynn

## Abstract

A major unresolved question in the origin and evolution of life is whether a continuous path from geochemical precursors to the majority of molecules in the biosphere can be reconstructed from modern day biochemistry. Here we simulated the emergence of ancient metabolic networks and identified a feasible path from simple geochemically plausible precursors (e.g., phosphate, sulfide, ammonia, simple carboxylic acids, and metals) using only known biochemical reactions and models of primitive coenzymes. We find that purine synthesis constitutes a bottleneck for metabolic expansion, and that non-autocatalytic phosphoryl coupling agents are necessary to enable expansion from geochemistry to modern metabolic networks. Our model predicts punctuated phases of metabolic evolution characterized by the emergence of small molecule coenzymes (e.g., ATP, NAD^+^, FAD). Early phases in the resulting expansion are associated with enzymes that are metal dependent and structurally symmetric, supporting models of early biochemical evolution. This expansion trajectory produces distinct hypotheses regarding the timing and mode of metabolic pathway evolution, including a late appearance of methane metabolisms and oxygenic photosynthesis consistent with the geochemical record. The concordance between biological and geological analysis suggests that this trajectory provides a plausible evolutionary history for the vast majority of core biochemistry.

## Introduction

Modern metabolism evolved as a consequence of nascent life deriving material and energy from the surrounding geochemical environment ^1–3^. However, the transition from geochemistry to extant biochemistry is poorly understood, due in part to a great uncertainty in the structure of ancient metabolic networks ^4^. In particular, chemical reactions that are unrelated to biochemistry have been invoked as missing steps in early biosynthetic pathways ^5–7^, suggesting that records of these chemical transformations were lost throughout the history of evolution; for example, through the emergence and sophistication of protein catalysts. Nevertheless, it is unclear to what degree ancient metabolism has been lost, and whether intermediate stages can be excavated from the extant biosphere.

Many lines of evidence suggest that recovering continuity between ancient geochemistry and extant biochemistry might be impossible without the inclusion of a vast number of abiotic chemical reactions unrelated to modern biology. First, a recent analysis of metabolic networks revealed a high prevalence of autocatalytic subnetworks in which the generation of several key biomolecules, including many coenzymes, are required for their own synthesis ^8,9^. Although autocatalysis may have been a necessary feature of early evolutionary processes ^8,10–12^, the widespread occurrence of such network motifs presents a problem for the initial emergence of ancient metabolism if there are no other routes for biosynthesis. Second, high rates of species extinction, horizontal gene transfer, non-orthogonal displacement, and evolutionary forces like drift could have eroded an early record of ancient biochemistry throughout the course of Earth’s history ^13–16^. Lastly, several recent studies simulating the emergence of metabolic networks from geochemistry only recovered small or fragmented networks, on the order of 10% of contemporary biochemistry ^17–20^. Likewise, models of ancient metabolism with hypothetical non-phosphate alternatives ^18,19^ drastically limited the potential coverage of modern day metabolic networks, which rely heavily on phosphate-containing molecules. Taken together, it remains unclear to what extent “extinct” biochemistry is necessary to enable the generation of modern metabolism from early Earth environments.

Beyond the question of continuity, constructing a model for the emergence of biochemical networks enables us to address key evolutionary questions. For example, resolving the order that biochemical reactions emerge in time can inform metabolic pathway evolution more broadly, including their relative ages and potential influence on geochemical and isotopic ‘biosignatures’ in the geologic record. Models of metabolic pathway evolution have invoked several mechanisms, ranging from sequential models, where reactions emerge in the order they appear in the reaction pathway, to mosaic models, where the order of reaction emergence is decoupled from the order of reactions in the extant pathway ^21–26 25,27^. Although studies have shown support for various models of evolution for specific pathways ^27^, a broad, biosphere-scale analysis of the relative occurrence of various modes of metabolic pathway evolution is lacking. Additionally, knowledge of the relative ordering of metabolic pathways that mediate biogeochemical cycling, such as carbon fixation^28^, can support efforts to interpret isotopic signatures in the geologic record.

Here, we construct a biosphere-level model of metabolic evolution and show that a single autocatalytic bottleneck in purine synthesis prevents the emergence of metabolism from geochemical precursors. We show that including a hypothetical ATP-independent pathway for purine biosynthesis enables the continuous expansion of metabolism from simple starting material, and that the ensuing trajectory of metabolic network evolution is correlated with features typically associated with the transition from ancient to modern biochemistry. We use this trajectory to resolve key aspects on the nature of metabolic evolution, with a focus on elucidating the mechanisms and order by which metabolic pathways emerged in the biosphere.

## Results

### Primitive purine production enables expansion to modern biochemistry

To construct a model of the evolutionary history of metabolism at the biosphere scale, we compiled a database of 12,263 biochemical reactions from the KEGG database (Table S1-3, Methods) ^29^. Unlike prior studies ^18,19^, we added detailed organic and inorganic cofactor dependencies for 5,259 reactions from UniProt, Expasy, PDBe, and EBI into the network (Table S1). These dependencies range from inorganic metal ions (e.g., Fe, Mn) to organic molecules (e.g., flavins, quinones) involved in catalysis (see Methods). Using this network, we performed network expansion (Methods, ^17–20,30–32)^ starting from a set of “seed compounds”. Our seed compounds included metals and inorganic material (e.g., Fe, Mn, Zn), CO_2_, hydrogen sulfide, molecular hydrogen, orthophosphate, ammonia, and 19 organic substrates that can be produced abiotically with iron, pyruvate, and glyoxylate. The choice of seed set compounds implicitly assumes that ancient carbon fixing reactions, similar to those found in the reductive tricarboxylic acid (rTCA) cycle and reductive acetyl-CoA pathway, are capable of producing simple carboxylic acids from CO_2_ and reductants like H_2_ ^33–38^ (Table S4 and see supplemental text on succinate semialdehyde). Consistent with previous studies ^18,19^, we could generate a network of 429 compounds from diverse pathways in central metabolism, including amino acid biosynthesis and some simple organic coenzymes like PLP (Fig. 1a, black line, Table S5). However, as our biosphere-level network consists of >8000 compounds, this scope only constitutes ∼5% of all known biochemicals, leaving the vast majority of molecules unreachable from simple seed compounds. Although the inclusion of primitive thioester energy coupling mechanisms and reductants may have been important during the early stages of biochemical evolution ^18,19^, these modifications marginally increased the scope of the expansion (*n*=734, Fig. 1d). Hypothesizing that our results were biased and limited by the inclusion of only cataloged biochemical reactions, we explored the possibility that unknown biochemical reactions could enable a more extensive expansion. To investigate this possibility, we included reactions from a database of hypothetical biochemistry ^39,40^, which added 20,183 new reactions to our network and increased the total size by a factor of ∼2.7. Repeating the expansion with this expanded reaction set resulted in only a slight increase in scope to 472 compounds (Extended Data Fig. 1), suggesting that neither currently cataloged nor predicted biochemistry contain transformations required to reach the vast majority of known metabolites.

**Fig. 1:**
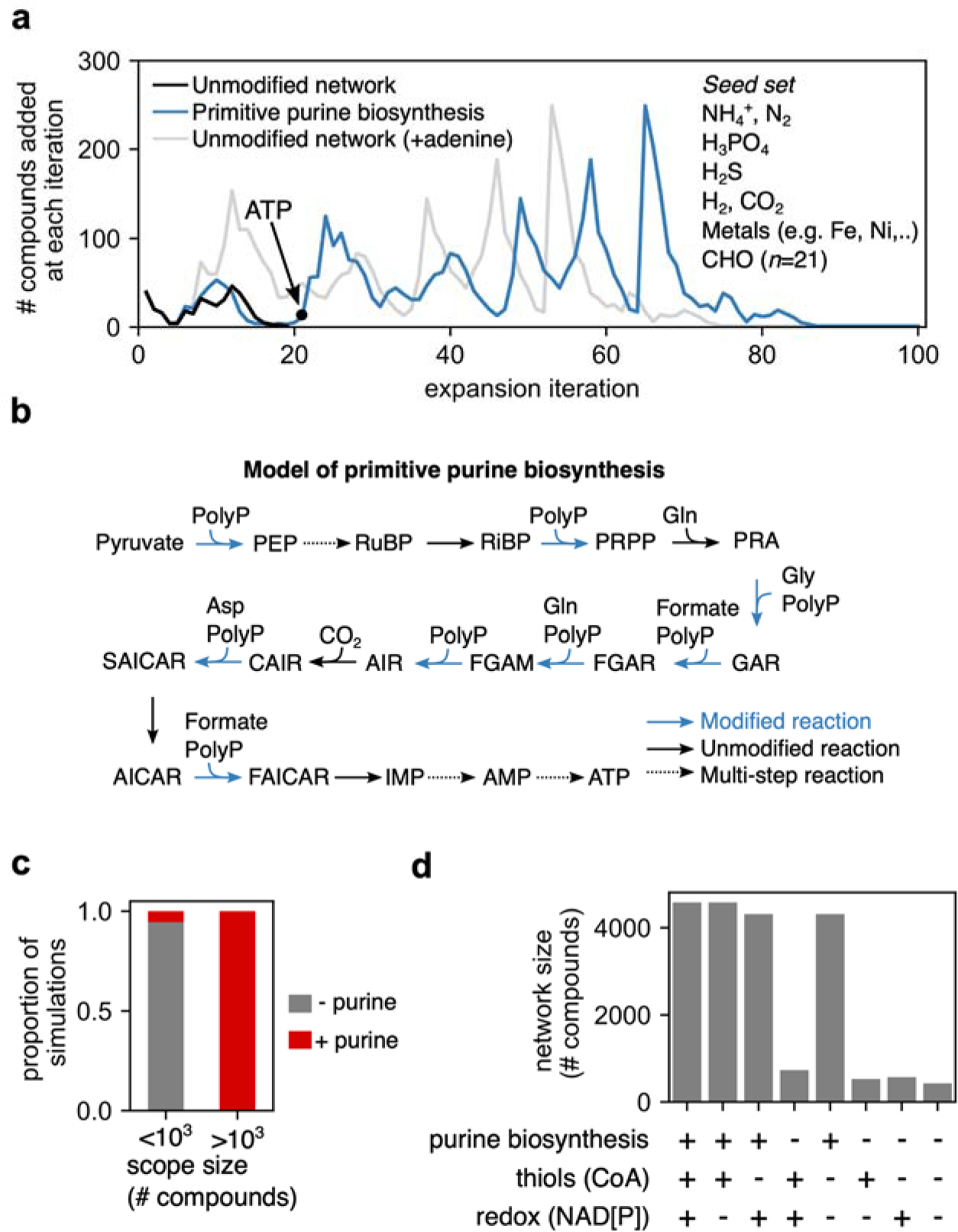
Primitive purine biosynthesis enables continuity between prebiotic seed molecules and contemporary biochemistry. (a) We performed network expansion with compounds previously hypothesized to have been highly abundant on early Earth (e.g., H_2_S, NH_4_^+^, CO_2_, Fe, Mn), as well as molecules producible in simple experiments from pyruvate, glyoxylate, and iron ^35^ (see Table S4 for complete list). For each simulation, we plotted trajectories of the number of compounds added (*y*-axis) at each expansion iteration (*x*-axis), with either an unmodified network without adenine (black line), with adenine (gray line) or a modified network with a model of a primitive polyphosphate (PolyP)-dependent purine biosynthesis pathway (blue line). The addition of adenine enabled the synthesis of ATP via the reversal of AMP:phosphate (5-phospho-alpha-D-ribosyl)transferase. (b) A model for primitive production of purines. We propose that all ATP-dependent steps in *de novo* purine biosynthesis were preceded by polyphosphate (PolyP) dependent steps (blue, ^43^). Full names for the abbreviations of these compounds are listed in the Materials and Methods. (c) For 8314 compounds, we added each as additional molecules in the seed set, repeated the expansion, and found that compounds containing purine moieties were required for expansion (see Methods). (d) We repeated expansion with models of primitive purine biosynthesis, primitive thioester coupling, or primitive redox systems (in lieu of NAD(P)). While networks increased slightly with primitive redox and thiol coupling, expansion to the >4000 compounds required primitive purine biosynthesis.

Notably, phosphoribosyl pyrophosphate (PRPP), a key precursor to metabolite classes like purines, was not in the expansion scope, suggesting that a bottleneck in purine production limits expansion. Indeed, the addition of adenine to the seed set resulted in a network of 4315 compounds, or ∼50% network coverage (Fig. 1a, gray line), including all major coenzymes (ATP, NAD, CoA, SAM, flavins, pterins, quinones, and heme; Table S5). To test whether purines were uniquely essential for the expansion to larger networks, we conducted a “rescue” experiment: for each of the 8314 single compounds from KEGG that were not produced in the original expansion scope (Fig. 1a, black line), we added one to the original seed set and repeated the expansion. We found that only 217 compounds yielded expansion to >4000 compounds, and all of these compounds either contained a purine moiety or were intermediates in *de novo* purine biosynthesis (Fig. 1c).

*De novo* purine production in extant biochemistry exhibits an autocatalytic dependence in the production of adenosine triphosphate (ATP) and PRPP, due to multiple steps requiring ATP as a phosphorylating agent (Fig. 1b)^8,9,41,42^. This autocatalytic dependence may have been relaxed in primitive metabolism, where primitive phosphorylating agents could have been either polyphosphate or acyl-phosphate (e.g., acetyl phosphate) ^43–45^. We explored this hypothesis by substituting 8 ATP-coupled reactions involved in purine synthesis with polyphosphate-coupled variants (Methods, Fig. 1b). We repeated the expansion of the original seed set with this modified network and found that expansion led to networks consisting of >4000 molecules with simple high-energy phosphoryl donors (Fig. 1a, blue line, Extended Data Fig. 2). To further explore the sensitivity of the expansion to variations in the seed set, we performed additional simulations where we first varied the composition of organic molecules, nitrogen, and sulfur sources and then re-ran the expansion algorithm. We found that expansion to networks with >4000 compounds required fully reduced nitrogen (e.g., ammonia or glycine) and relatively oxidized carbon source (Extended Data Fig. 3). Taken together, this model demonstrates that expansion from reduced nitrogen and oxidized carbon to most of primary metabolism requires only a small number of additional phosphate-coupled reactions involved in *de novo* purine biosynthesis.

### Punctuated metabolic innovation is gated by coenzyme synthesis

Along the primitive purine biosynthesis expansion trajectory (Fig. 1a, blue line), compound addition occurs in bursts, representing periods where large numbers of metabolites were added simultaneously. This observation motivated us to explore the hypothesis that production of key intermediate molecules are bottlenecks throughout the expansion. To test this hypothesis, we removed a single metabolite (and all associated reactions) from the network, performed network expansion on the perturbed network, and then repeated this procedure for each metabolite in the expansion scope (Fig. 2a). Of the 4,248 compounds produced during the expansion, only the removal of 149 led to scopes less than 4000 compounds, indicating that the vast majority of perturbations had little effect on the network size. By plotting the size of the network perturbed by removing a compound (*y*-axis) versus the iteration where the compound was made in the original expansion (*x*-axis), we found that removing certain compounds produced networks of similar size to networks generated before the compounds were produced in the unperturbed expansion (Fig. 2a, red dots). This result suggests that these compounds represent a specific class of “bottleneck” molecules that restrict expansion and are made exactly when they are essential for further expansion of the network (Fig. 2a, red dots). Interestingly, several of these molecules were coenzymes like PLP, ATP, NAD, CoA, and thiamine diphosphate (ThDP), or intermediates in the biosynthesis of key coenzyme classes like isoprenoids (isopentenyl diphosphate).

**Fig. 2:**
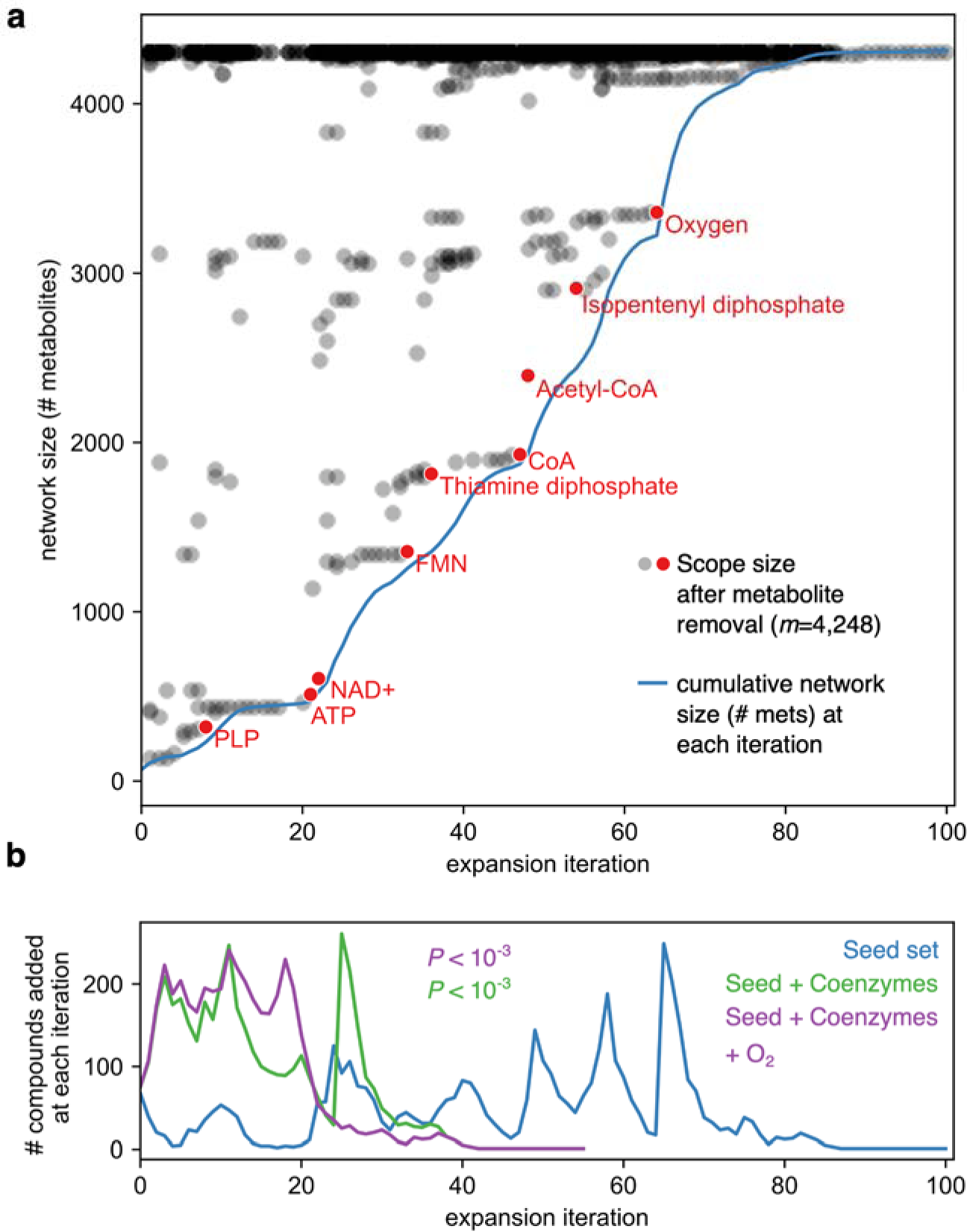
Peaks during metabolic expansion are associated with coenzyme and oxygen production. (a) For each compound in the expansion scope (*m*=4,248), we removed the compound and all associated reactions from the network used in Fig. 1 and performed expansion. For each compound, we plotted the iteration when the compound is formed in the original expansion (*x*-axis) versus the scope size achieved in the perturbed expansion (*y*-axis). As a reference, we also plotted the cumulative network size (number of compounds) after each iteration for the unperturbed expansion (blue line). Red points are examples of important compounds that become limiting for the expansion close to when they are produced, suggesting they are bottlenecks in the expansion. (b) Trajectories of compounds added at each iteration during network expansion with the base seed set (*n*=70, blue line), base seed set with eight additional coenzymes (ATP, NAD, CoA, PLP, FAD, SAM, ThDP, and Cobalamin), (*n*=78, green line), and the base seed set with additional coenzymes and molecular oxygen (*n*=79, purple line). We measured the iteration where 90% of the network was recovered, and found that this value was significantly less than simulations with eight randomly selected compounds as additional seed molecules (Monte Carlo permutation test: *P* < 10^-3^).

As noted in previous studies, the inclusion of various coenzyme classes has a large effect on metabolic network expansion ^17,30^, motivating the hypothesis that the observed peaks were induced by the production of key metabolites. To explore this possibility, we simultaneously added 8 key coenzymes (ATP, NAD, CoA, PLP, FAD, SAM, ThDP, and cobalamin) as seed molecules and re-ran the expansion. Including these coenzymes collapsed the multiple peaks observed in Fig. 1a into just two peaks, where the later peak corresponds to the production of molecular oxygen (Fig. 2b). We further found that addition of these 8 coenzymes to the seed set enabled expansions with fewer iterations relative to randomly chosen sets of 8 compounds with or without oxygen (Monte Carlo permutation test: *P* < 10^-3^). Altogether, this analysis shows that the production of major coenzymes are key factors shaping the observed peaks during expansion.

### Ancient features of compounds and reactions are correlated with expansion iteration

The last peak shown in Fig. 1a corresponds to the production of molecular oxygen after the emergence of quinones used in modern-day photosystems (Supplemental Text). Since O_2_ production was only thought to emerge ∼2.4 Gya, after the rise of cyanobacteria ^46,47^, it is tempting to hypothesize that expansion iteration number may, to a first approximation, represent time. To investigate this possibility, we computed metrics associated with ancient biochemistry and examined how these metrics changed throughout the expansion process.

As suggested recently ^48^, the complexity of molecules (as well as their abundance) might be a useful metric to quantify the degree of evolution of the biosphere. To explore this possibility, we first computed the Bertz complexity of each molecule with a defined molecular structure that was produced during the expansion (*n*=3588, Table S6). For each iteration of the expansion, we identified all the compounds produced at that iteration and computed the mean Bertz molecular complexity. The expansion iteration number was positively correlated with mean molecular complexity (Fig. 3a, Pearson’s *r* = 0.72, *P* < 10^-16^) consistent with the hypothesis of increasing molecular complexity during metabolic evolution. This trend was also observed with other measures of molecular complexity, and large expansions were only observable with relatively simple seed sets compared to more complex seed sets (Extended Data Fig. 4). The mean molecular complexity is correlated with expansion iteration even when controlling for the average redox state of molecular carbon (Partial correlation: *r*_partial_ = 0.70, *P* < 10^-15^).

**Fig. 3:**
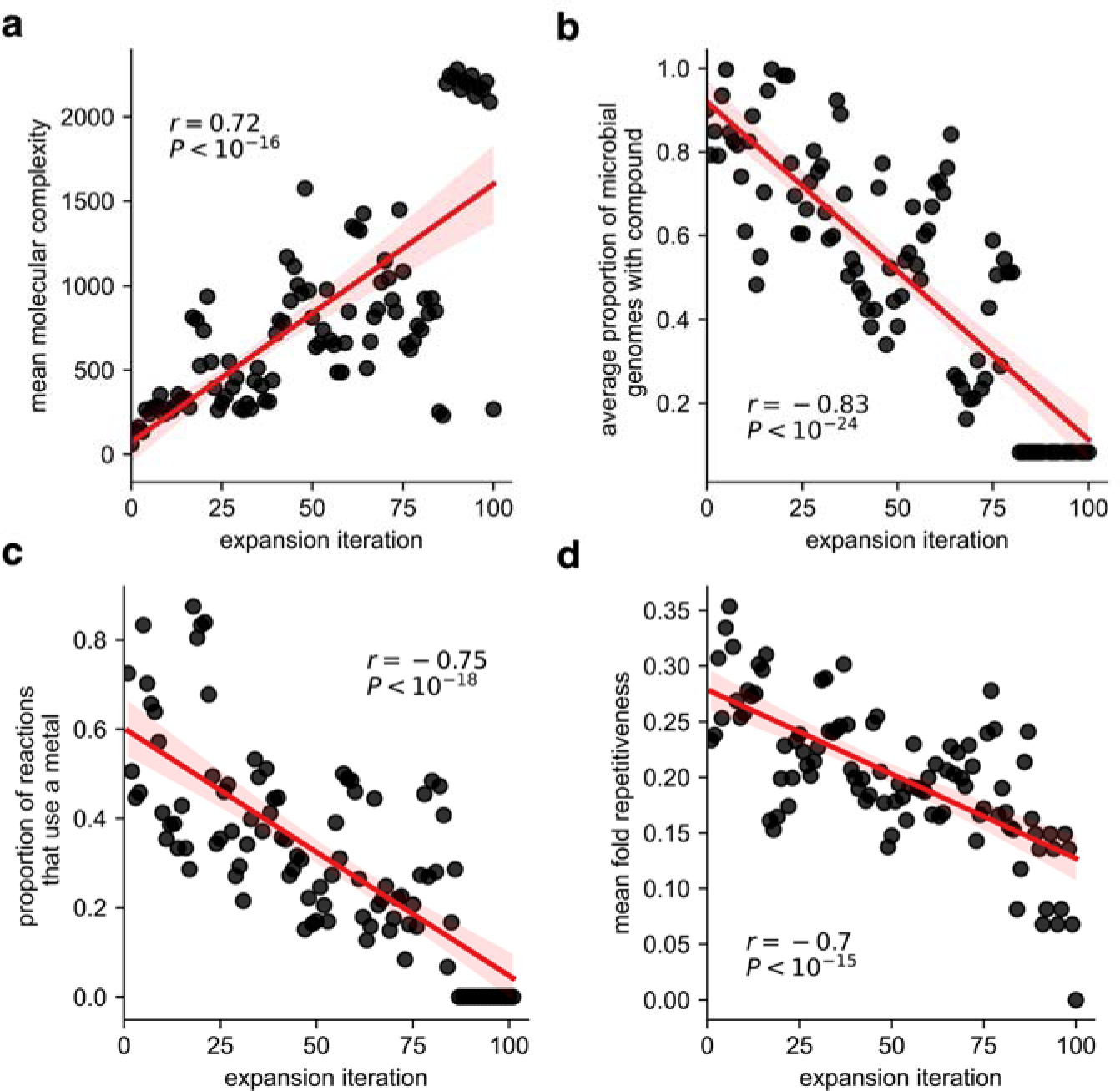
Ancient biochemical features are correlated with expansion iteration. We computed various biochemical features previously hypothesized to be associated with ancient biochemistry at each iteration during the expansion (*x*-axis, a-d). We computed (a) the average Bertz molecular complexity of molecules produced at each expansion step (*y*-axis), (b) the average proportion of microbial genomes using each compound produced during each expansion step (*y*-axis), (c) the proportion of reactions that use an enzyme dependent on a metal cofactor (*y*-axis), and (d) the mean repetitiveness for protein folds associated with each reaction utilized at each expansion iteration (*y*-axis) (see Methods).

The proportion of extant prokaryotic genomes using compounds produced during each expansion step decreased throughout the expansion (Fig. 3b, Pearson’s *r* = −0.83, *P* < 10^-24^, see Methods), consistent with the hypothesis that the most ancient parts of metabolism are the most conserved across the tree of life. More broadly, we found that this network is biased for compounds in pathways that are frequently observed in prokaryotic genomes and, on average, compounds in this network are found in 55% of prokaryotic genomes, compared to just 10% for KEGG compounds not included in the expansion scope (Extended Data Fig. 5). At each iteration, we computed the proportion of reactions that are exclusively found in eukaryotic genomes (see Methods). Interestingly, the proportion of reactions unique to eukaryotes increases towards the end of the expansion (Extended Data Fig. 6a). The selective removal of eukaryote-specific reactions (*n*=1638) led to a network of 3806 compounds (Extended Data Fig. 6b), which is larger than expansions performed after the removal of an equal number of randomly selected reactions (Monte Carlo Permutation test: *P*<10^-3^).

Models of ancient metabolic evolution suggest that before the evolution of the modern genetic coding system, “pre-enzymatic” catalysts such as minerals, metal ions, peptides or short RNA polymers served as catalysts for the majority of metabolic reactions ^34,49–52^. It has been proposed that relics of these pre-enzymatic catalysts may be retained in the active sites of many extant enzymes as metal or iron-sulfur cofactors ^4,33,53^. As Archean oceans became increasingly sulfidic and/or oxidized ^54^, the geochemical supply of various metals like iron may have decreased. This decline in the solubility of iron-bearing minerals may be reflected in the increased deposition of iron formations after ∼3 billion years ago ^55^ and may have provided a selective pressure for new enzymes to be less dependent on specific metals and iron-sulfur cofactors. We thus computed the proportion of reactions at each iteration that were metal cofactor-dependent (metal cofactors are listed in Table S8), and observed that this proportion generally decreased as expansion iteration increased (Fig. 3c, Pearson’s *r* = −0.75, *P* < 10^-18^), consistent with the idea that later networks were less dependent on metal-catalyzed reactions.

Beyond molecular complexity, taxonomic associations, and metal-dependence in reaction mechanisms, specific types of protein folds have been hypothesized to have served as ancient catalysts in primitive biochemistry ^56–59^. In particular, it has been argued that repetitive/symmetric folds may have been among the first protein folds used in metabolism ^60^, due to their ability to encode complex structures within a short, oligomerizing peptide ^61–64^ and their ubiquitous association with diverse metabolic processes ^56,65,66^. Assuming ancient protein folds had more opportunity to become catalysts for biochemical reactions than more recent protein folds, we hypothesized that iteration number should correlate with enzyme fold repetitiveness, a putative correlate of fold age. To explore this possibility, we computed a structural repetitiveness score for the set of protein folds associated with a reaction ^67^, and the mean repetitiveness score at each expansion iteration (Table S9, Methods). Consistent with our expectation, we observed a significant decrease in mean repetitiveness throughout the expansion trajectory (Fig. 3d, Pearson’s *r* = −0.70, *P* < 10^-15^). This association is still observed when limiting our analysis to reactions dependent on a single enzyme family (Pearson’s *r* = −0.49, *P* < 10^-6^). An alternative explanation for this trend is the potential association between enzyme and substrate complexity. We thus computed the partial correlation between structural repetitiveness and expansion iteration, controlling for molecular complexity, and still observed a strong decrease in repetitiveness throughout the expansion trajectory (Pearson’s partial correlation *r*_partial_ = −0.72, *P* < 10^-15^).

Altogether, these analyses show that many features of ancient metabolism – including molecular complexity metrics, metal-cofactor dependencies, and protein fold attributes – are associated with expansion iteration, consistent with the hypothesis that expansion iteration can be viewed as a first-order approximation of time along the trajectory of metabolic evolution.

### The modes and order of emergence of extant biochemical pathways

The trajectory of biochemical network evolution permitted us to explore fundamental questions in the history of metabolism, including how metabolic pathways evolve and the specific ordering of distinct metabolic pathways. We first sought to determine the degree to which extant linear metabolic pathways evolved sequentially (either in the forward or reverse direction, see Methods), or non-consecutively, where pathway steps are realized at various times, independent of pathway structure (i.e., the mosaic model ^22–24)^. Sequential models of metabolic evolution include the forward direction (also called the Granick, or prograde model) ^68^, where reactions emerge following the proceeding reaction in the metabolic pathway, and the reverse direction, where reactions are added sequentially in a “retrograde” fashion ^69^. For the set of linear metabolic pathways in the KEGG database reachable after expansion (*n*=110, Table S10), we computed the iteration where each pathway step was feasible and classified these pathways based on the Spearman’s rank correlation between iteration number and when each step was feasible (Methods, Fig. 4a). While we found that that ∼13% (15/110) of pathways adhere to a retrograde model of pathway evolution, including core pathways like cysteine biosynthesis from serine (Table S10), and ∼49% (54/110) follow a prograde model, a large fraction (41/110) of these pathways do not emerge sequentially, but rather emerge in a discontinuous fashion, consistent with a mosaic model of metabolic evolution.

**Fig. 4:**
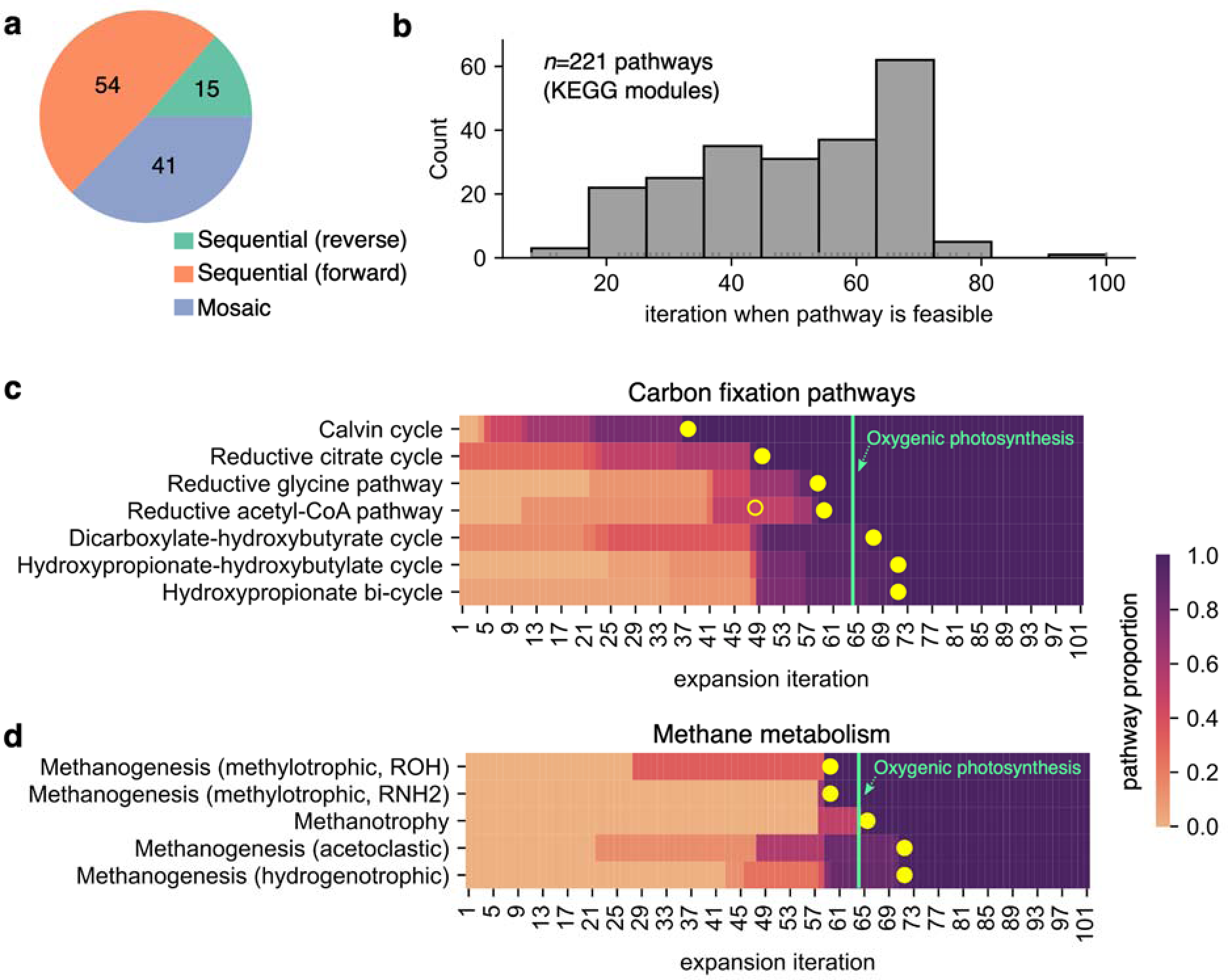
Timing the emergence of metabolic pathways. (a) For 110 linear metabolic pathways in the KEGG database (Table S10), we classified each pathway as emerging sequentially (either via the forward or reverse direction) or non-sequentially (i.e., Mosaic, blue). We found that 37% (41/110) of these pathways emerged non-sequentially and ∼13% (15/110) adhere to a retrograde model of pathway evolution. (b) For 221 KEGG modules, we computed and plotted the distribution of the minimum expansion iteration when each module became feasible (Table S11). (c) A heatmap showing the proportion of feasible steps (color) of seven carbon fixation pathways (*y-*axis) throughout the expansion (*x*-axis). The closed yellow dots represent the iteration when all steps in the pathway are feasible. The open yellow circle denotes the emergence of the carbonyl-branch of the reductive acetyl-CoA pathway. (d) The same as (c) but for pathways involved in methane metabolism. The green vertical line in both subplots (c) and (d) denote the iteration when oxygenic photosynthesis emerged.

We next sought to order the emergence of specific extant metabolic pathways. We constructed logical relationships between reactions and pathways accounting for redundancies and co-dependencies (see Methods), and computed the iteration number where each of 221 KEGG modules were feasible (Table S11, see Methods). Pathway emergence is distributed throughout the entire expansion (Fig. 4b). For example, before ATP is generated in the network, only the reductive and non-oxidative pentose phosphate modules, PLP biosynthesis, and cysteine biosynthesis are fully enabled (Table S11). In order to make specific predictions testable with the geologic record, we analyzed the ordering of pathways involved in key biogeochemical cycling in the biosphere, such as carbon fixation and methane metabolism. Surprisingly, of the seven known carbon fixation pathways, the Calvin cycle was predicted to become feasible first, preceding both the reductive tricarboxylic acid cycle (rTCA) cycle and the reductive acetyl-CoA pathway, which have both been predicted to be operative in the last universal common ancestor (Fig. 4c) ^70–72^. While the relative order between the rTCA and Calvin cycle was dependent on the seed set composition, we found that the extant Calvin cycle emerged before the extant reductive acetyl-CoA pathway in all simulations with random seed sets (Extended Data Fig. 7). Consistent with this observation, we found that pathways involved in archaeal methane metabolism also emerged after the Calvin cycle and rTCA cycle (Fig. 4d), due to the heavy reliance on pterin cofactors. While methylotrophic and hydrogenotrophic variants of methanogenesis have both been proposed as being ancestral in ancient archaea ^73^, network expansion predicts that methylotrophic methanogenesis preceded the hydrogenotrophic variant. Fractionations of ^13^C from ^12^C in 2.9-2.7 billion year old sediments ^74^ are consistent with the operation of methanogenesis ^75^ and/or acetogenesis ^76^, both of which rely on the reductive acetyl-CoA pathway today. However, the global distribution of older samples (from 3.0 to 4.1 billion years) display C-isotope fractionations akin to those generated by the modern day Calvin cycle ^74^. While we cannot rule out alternative metabolic explanations for these isotopic patterns (see Supplemental Note), our metabolic network evolution model, like these geological data, suggests the emergence of the Calvin cycle preceding the modern variant of the reductive acetyl-CoA pathway.

If the Calvin cycle developed early in biosphere evolution, an enzyme with ribulose-1,5-bisphosphate carboxylase (RuBisCo) activity must also have appeared early. While Form I RuBisCo is confined to cyanobacteria, eukaryotic algae, and some proteobacteria, other RuBisCo forms (e.g., Forms II, II/III, III) and RuBisCo-like proteins (RLPs) are more widespread and possibly ancestral ^77^, suggesting that the earliest variants of the Calvin cycle could have used these variants for carboxylation.

## Discussion

Continuity between early Earth geochemistry and modern metabolism is a necessary condition for any theory of the emergence and evolution of the biosphere on Earth. Whereas demonstrating a continuous trajectory does not rule out possibly critical, yet extinct chemistry (for example ^5,6^), it does relax the requirement to invoke such transformations as essential for biochemical evolution. Furthermore, whereas many other feasible trajectories may enable this observed continuity, the trajectory in Fig. 1a recapitulates several features associated with time and Earth history. Our model suggests that, while species extinction is likely ubiquitous throughout time, extinction of primary metabolic biochemistry at the biosphere scale may in fact be quite rare. With an approximate time ordering of metabolic exploration, we can explicitly probe the ordering of extant metabolic pathways and key biochemical innovations in the biosphere^78,79^. It is important to note, however, that our model can only predict the timing of extant biochemical pathways and not necessarily their primitive analogues. Our analysis suggests that the evolution of the extant rTCA cycle may have been limited by the emergence of cofactors like CoA and thiamine pyrophosphate. Importantly, our choice of seed compounds suggest that ancient analogues of carbon fixation pathways similar to the rTCA and reductive acetyl-CoA pathways produced many important carboxylic acids, as explored in previous theoretical and experimental work^18,19,25,35–38,71^.

The peaks of metabolic expansion may be characteristic of evolving metabolic networks. Indeed, we found that this punctuated structure is robustly observed when perturbing the initial seed set composition (Extended Data Fig. 4), when we include a database of predicted extant yet undiscovered biochemistry (Extended Data Fig. 1), and when we remove eukaryote-specific biochemical reactions (Extended Data Fig. 6). However, the particular structure and trajectory in Fig. 1a may be sensitive to seed set molecule composition or network structure (Fig. 2b, Extended Data Fig. 3,4 and 7). In contrast, the emergence of metabolic pathways appears to occur with minimal punctuated structure (Fig. 4b), highlighting key differences between the shape of metabolic evolution at the scale of individual reactions and compounds versus pathways. A large fraction of metabolic pathways also appear to occur as a coalescence of biochemistry emerging at different times throughout the expansion, suggesting the “mosaic” model of metabolic evolution might be a major mode of evolution. The core metabolic pathways in extant species might have emerged relatively late, with primitive metabolic functions enabled instead by biochemical transformations from disparate metabolic pathways, like previously proposed ancient carbon fixation pathways ^19,71 34,71,80^. Moreover, it has been suggested that modern biochemical pathways may contain relics of such an evolutionary model, due to the repetitiveness of their chemical reaction sequences ^81^.

Although improvements in biochemical annotation and thermodynamic parameter estimation will undoubtedly change details of the trajectory presented in this paper (see Supplemental Information), our analysis suggests that adding an ATP-independent purine biosynthesis pathway to the known catalog of extant biochemistry is necessary to enable the continuous trajectory from geochemical precursors to the majority of core metabolism. Efforts to model metabolic network evolution from geochemically-plausible molecules were unable to generate purines, despite extensive inclusion of non-catalogued hypothetical biochemistry like thioester-coupling ^18,19^. While abiotic synthetic routes to purines are well studied both experimentally ^82,83^ our analysis suggests that these abiotic mechanisms may have not been necessary if analogues to modern-day phosphate-coupling mechanisms were accessible during the development of the biosphere ^44^.

Prior studies have uncovered several autocatalytic compounds enriched in autocatalytic sets at both organismal and biosphere scales ^8,9,84^, including ATP, PLP, and NAD. Contrary to what one might expect from these studies, we find purine biosynthesis is the only autocatalytic pathway that limits expansion of biosphere-scale metabolism. Compared to the wide occurrence of strict autocatalytic dependencies at the genome-level ^42,84^, there appears to be a paucity of obligate autocatalytic dependencies in the biosphere-level metabolism. This observation is consistent with the hypothesis that autocatalytic pathways are strongly selected for at the organismal scale ^85,86^, yet primitive, non-autocatalytic pathways can be reconstructed by analyzing biochemistry at higher scales of biological organization. These results suggest that biosphere-level metabolic networks, unlike organismal-scale metabolism, may be the appropriate scale to interrogate evolutionary history of metabolism in deep time^8,17,18,32,80^. An intriguing possibility is that this single autocatalytic dependency observed here is due to the incomplete characterization of extant biochemical diversity, and that fragments of an ancient, non-autocatalytic polyphosphate-dependent *de novo* purine synthesis pathway remains hidden in the biosphere.

## Acknowledgements

The authors thank Woodward Fisher, Avi Flamholz, and Joan Valentine for valuable discussions. J.E.G. is supported by the Gordon and Betty Moore Foundation as Physics of Living Systems Fellows through grant number GBMF4513, as well as the Simons Foundation. S.E.M. and B.A.W. acknowledge support from NSF Awards 1724300 and 1724393(Collaborative Research: Biochemical, Genetic, Metabolic, and Isotopic Constraints on an Ancient Thiobiosphere). S.E.M. additionally acknowledges support from JSPS KAKENHI (Grant Nos. JP18H01325 and 22H01343). H.B.S acknowledges support from JSPS KAKENHI Grant No. JP19K23459.

## Contributions

All authors designed the research. J.E.G, H.B.S. and L.M.L. prepared data. J.E.G. and H.B.S. wrote code and ran simulations. J.E.G. performed analysis. J.E.G., H.B.S. and L.M.L wrote the manuscript. All authors read and approved the final manuscript.

## Materials and Methods

### Software availability

Code and data are available on the following github repository: https://github.com/jgoldford/metabolic-continuity

### Metabolic network reconstruction

Briefly, we downloaded the KEGG network on May 31, 2021., We removed *n*=230 reactions that were elementally inconsistent (i.e., the reactants and products did not share the same elements) (Table S3). Reactions that were stoichiometrically imbalanced, or relied on metabolites with unknown “R-groups”, were kept. All reactions are listed in Table S1.

#### Missing reaction and cofactor annotation

Cofactors and EC-cofactor dependencies were identified from Expasy (via the ‘cofactorLabel’ field and comments), PDBe (via the cofactor summary), UniProt (via cofactor/EC numbers), and KEGG (via comments and reaction definitions). For more details, see the SI section titled “Cofactor Annotations”. We added several missing reactions in KEGG to ensure production of key, generic metabolites, such as quinones, ferricytochrome. We describe each modification in the SI section titled “Addition of missing reactions.”

#### Thermodynamic constraints

Standard molar free energies were estimated by the component contribution method ^87^ and the eQuilibrator python API ^88^ using the following parameters: pH=7.0, pMg=3.0, ionic strength of 0.25M and *T*=298.15. For each reaction with a free energy estimate at standard molar conditions, we estimated the maximum and minimum reaction free energy for both the forward and reverse reactions. We computed these values by assuming that compound concentration ranges were between 10^-7^ and 10^-1^ M, and computed the maximum and minimum reaction free energies as described previously ^19^. For all calculations, we assumed activity coefficients were well approximated by concentrations. All forward or reverse reactions with minimum free energies above zero were removed from the network. For KEGG reactions with no free energy estimate (*n*=839), we kept both the forward and reverse reaction in the network.

#### Modeling primitive purine biosynthesis reactions

We modeled the primitive purine biosynthesis by adding 8 new reactions to the model. We replaced all ATP (GTP) with pyrophosphate (PP_i_) (C00013) and ADP (GDP) with orthophosphate (C00009) and recomputed standard molar free energies using eQuilibrator for the following KEGG reactions: 5-Phospho-D-ribosylamine:glycine ligase (R04144, EC 6.3.4.13), ATP:pyruvate 2-O-phosphotransferase (R00200, EC 2.7.1.40), formate:5-amino-1-(5-phospho-D-ribosyl)imidazole-4-carboxamide ligase (R06975, EC 6.3.4.23), ATP:ribose-1,5-bisphosphate phosphotransferase (R06836, EC 2.7.4.23), 1-(5-Phosphoribosyl)-5-amino-4-carboxyimidazole:L-aspartate ligase (R04591, EC 6.3.2.6), 5’-Phosphoribosylformylglycinamide:L-glutamine amido-ligase (R04463, EC 6.3.5.3), formate:N1-(5-phospho-beta-D-ribosyl)glycinamide ligase (R06974, EC 6.3.1.21), 2-(Formamido)-N1-(5-phosphoribosyl)acetamidine cyclo-ligase (R04208, EC 6.3.3.1). In Fig. 1b, we provide a plausible pathway with just 8 of these reactions. The abbreviations for the compounds in this pathway are as follows: Gly = Glycine; Gln = L-Glutamine; Asp = L-Aspartate; PolyP = Polyphosphate (Pyrophosphate); PEP = Phosphoenolpyruvate; RuBP = D-Ribulose 1,5-bisphosphate; RiBP = D-Ribose 1,5-bisphosphate; PRPP = 5-Phospho-alpha-D-ribose 1-diphosphate; PRA = 5-Phosphoribosylamine; GAR = 5’-Phosphoribosylglycinamide; FGAR = 5’-Phosphoribosyl-N-formylglycinamide; FGAM = 5’-Phosphoribosyl-N-formylglycinamidine; AIR = Aminoimidazole ribotide; CAIR = 1-(5-Phospho-D-ribosyl)-5-amino-4-imidazolecarboxylate; SAICAR = 1-(5’-Phosphoribosyl)-5-amino-4-(N-succinocarboxamide)-imidazole; AICAR = 1-(5’-Phosphoribosyl)-5-amino-4-imidazolecarboxamide; FAICAR = 1-(5’-Phosphoribosyl)-5-formamido-4-imidazolecarboxamide; IMP = Inosine monophosphate; AMP = Adenine monophosphate; ATP = Adenine triphosphate.

#### Models of primitive thioester and redox coenzymes

Previous work suggested that primitive versions of CoA-coupled and NAD(P)-coupled reactions could have enabled the generation of metabolic networks before the availability of phosphate ^18,19^. We explored whether primitive coenzyme systems could enable purine production by (i) substituting all CoA-coupled reactions with the thiol mercaptopyruvate and (ii) substituting all NAD(P)H-coupled reactions with hydrogen gas. Note for (i), we were unable to re-compute standard molar free energies because the thioesters formed from mercaptopyruvate were not in the KEGG database. For (ii), we computed standard molar free energies as described previously.

#### Extending KEGG with additional reactions from ATLAS

We obtained a list of all reactions in the ATLAS database ^39,40^. We removed reactions that matched KEGG reactions, resulting in an additional 20,183 reactions. Note that the ATLAS database also contained an additional 3024 metabolites, increasing the total network to 32,449 reactions and 11,768 metabolites. We applied the same thermodynamic methods described above for ATLAS reactions.

### Seed set

All seed molecules are listed in Table S4. The seed set consisted of all inorganic divalent and monovalent metal species in KEGG, as well as elemental sources of phosphorus, nitrogen, sulfur, oxygen, hydrogen, and carbon. While ammonia, orthophosphate and hydrogen sulfide were the only nitrogen, phosphorus, and sulfur sources provided in the seed set, respectively, several organic compounds were included as seed molecules. Since we did not assume a primitive thioester coupling mechanism and include thiols in the seed set, organic molecules such as pyruvate were necessary to enable expansion ^19^. We thus included organic molecules producible when reacting glyoxylate with pyruvate ^35^, resulting in 19 organic compounds (Table S4). Note that when we assumed that succinate semialdehyde could be producible from succinate via geochemically available reductants (e.g., H_2_), we were able to generate networks >4000 compounds with only pyruvate as the organic seed molecule.

### Network expansion algorithm

The network expansion algorithm was run as described previously ^18,19^. Briefly, seed compounds are allowed to react given the reactions in the network, which then produce product compounds. The product compounds are added to the seed set, and the process is repeated until convergence. The network expansion algorithm was implemented using the `networkExpansionPy` python package (https://github.com/jgoldford/networkExpansionPy).

### Identification of purine-containing molecules

Purine-containing compounds were identified using the SUBCOMP search function hosted by the KEGG database (https://www.genome.jp/tools/subcomp/). The query structure was purine (i.e., KEGG compound C15587), the database was “COMPOUND”, and the search mode was “SUBstructure”.

### Defining metabolic pathways

We defined metabolic pathways using KEGG modules (https://rest.kegg.jp/list/module). The ordering of reactions and reaction definitions for each module were determined from the “Reaction” field and “Diagram” field in each KEGG module entry. Data was retrieved using the TogoWS REST service via Biopython (https://biopython.org/docs/latest/api/Bio.TogoWS.html). Because the complexity and non-linearity of some modules make it hard to interpret which reactions are strictly necessary, we limit our analysis to linear pathways (Table S10). A full list of the KEGG modules included for analysis can be found in Table S11. Pathways containing less than 2 reactions are not included. For the linear pathways, we used the reaction to module step rules to compute when each step in the pathway was feasible during the expansion. We then computed the Spearman’s rank correlation between iteration number and pathway step (indexed as 0 to *N*, where *N* is the total number of steps in a pathway). A Spearman’s rank correlation of 1 indicated the pathway emerged sequentially in the forward direction, while a Spearman’s rank correlation of −1 indicated the pathway emerged sequentially in the reverse direction.

### Molecular complexity and other ancient features

To compute the molecular complexity for each compound in the expansion scope with a defined molecular formula (*n*=3588), we used RDKit (version 2022.03.1) (https://www.rdkit.org/) to compute the Bertz complexity ^89^ using the `GraphDescriptors` module with default parameters (Table S6). We computed topological indexes (Extended Data Fig. 6a-b), using custom python scripts, dependent on NetworkX (version 2.5.1). We computed the average state of reduction per carbon ^2^ (see Extended Data Fig. 3c, Table S7) on all organic molecules with only carbon, oxygen, and hydrogen atoms (*n*=1178).

We computed the proportion of microbial taxa using each compound in the expansion by using the KEGG REST API to download KEGG modules used in each prokaryotic genome (*n*=6416), as well as all KEGG compounds used in each KEGG module. We took 1,314 KEGG compounds that are intermediates in 377 KEGG modules, and computed the proportion of prokaryotic genomes that use each compound as an intermediate in at least one KEGG module. For all compounds produced and a specific expansion iteration, we computed the average proportion of genomes using each compound (Fig. 3b). For Extended Data Fig. 3, we identified KEGG reactions that were unique to Eukaryotes and Prokaryotes. We first identified enzyme classes (EC numbers) in 8652 genomes, 838 of which were Eukaryotes, and computed the EC classes that were unique to either Prokaryotic or Eukaryotic-specific. We converted these KEGG reactions, using KEGG’s internal mapping between reactions and EC number. We identified 1638 reactions that were unique to Eukaryotes.

### Calculating Reaction-Fold Associations

KEGG orthologous (KO) groups, 8030 in total, were clustered at 80% sequence identity by CD-HIT ^90^ using the default settings and a word size of 5 characters. Using the curated hidden Markov models (HMMs) from the Evolutionary Classification of Domains (ECOD) database ^91,92^ (ECOD version develop279), protein families were mapped onto the representative KEGG genes of each orthologous group by the program hmmsearch ^93^. HMM searches used the default settings and the search space (the -Z flag) was set to 106,052,079 sequences. Only domains with an independent E-value of less than 1×10^-4^ and an HMM coverage of greater than 70% were considered potential hits. Domains were assigned to regions of a gene in order of increasing E-value using the “envelope coordinates” of the associated hit. If two hits overlap by more than 30% of either hit, the hit with the higher E-value is discarded. Finally, reconciled hits associated with a gene were mapped to their corresponding ECOD X-groups when such mappings are unambiguous. An association between an X-group and KO group was identified if at least 20% of the genes contained that X-group domain.

We next constructed a mapping between reactions and KO groups. We used the KEGG REST API to download the associations between KEGG reactions and Enzyme Commission (EC) numbers, and the EC numbers and KEGG orthologous groups. A KEGG orthologous group was associated with a KEGG reaction if they were associated with at least one shared EC number. Finally, X-group and reactions associations were obtained via their mutual associations with KO groups.

### Computing structural repetitiveness of protein folds

Protein fold repetitiveness for each representative ECOD domain (70% sequence identity cutoff, ECOD version develop279) was calculated using CE-Symm 2.0 ^67^ with default settings. All classifications other than C1 (i.e., no internal symmetry) were defined as repetitive. Repetitiveness scores were taken to be the fraction of repetitive representative domains within a fold, defined here as an ECOD X-group (Table S9). The average repetitiveness score for a reaction was calculated by averaging the repetitiveness scores of all associated folds (Table S9). The average fold repetitiveness per expansion iteration (Fig. 3d) was computed by averaging across all reactions that emerged at each iteration step.

### Statistical analysis

Pearson’s correlation, Spearman’s correlations and two-sided, nonparametric Mann Whitney *U* tests were all performed using the `pearsonr`, `spearmanr` and `mannwhitneyu` functions from the `scipy.stats` python module, version 1.5.4 in python 3.6. For Monte Carlo permutation tests performed in Fig. 2b, we used the `random` function in the python native library `sample` to randomly sample 10^3^ combinations of 8 molecules from the expansion scope (but not the seed set in Table S4), and then repeated the expansion. We estimated a *p*-value by computing the proportion of randomly chosen seed sets that led to an expansion with fewer expansion iterations than the original expansion shown in Fig. 2b (blue line). Partial correlation analysis was performed using the `penguin.partial` (version 0.3.12) function in python.

