## Supplementary Information for "Primitive purine biosynthesis connects ancient geochemistry to modern metabolism"

### Supplemental Information for: **Primitive purine biosynthesis connects ancient geochemistry to modern metabolism**

#### Network reconstruction

In this supplemental note, we detail the network modifications made manually to account for missing cofactor annotations and gaps in the biosphere-level metabolic network reconstruction.

#### Cofactor annotations

Cofactors and EC-cofactor dependencies were identified from Expasy, PDBe, UniProt, and KEGG. Expasy data was accessed via the FTP (https://ftp.expasy.org/databases/enzyme/). For each entry in the `enzyme.rdf` file, the associated EC was identified via the `rdf:about` field, and associated cofactors were identified via the `cofactorLabel` field. Comments from the `rdfs:comment` field were preserved. PDBe data was accessed via the API (https://www.ebi.ac.uk/pdbe/graph-api/pdbe_doc/#api-Compounds-GetCofactorSummary / https://www.ebi.ac.uk/pdbe/graph-api/compound/cofactors/). Each entry in the JSON is a cofactor, and each EC in the subfield `EC` identifies ECs where that cofactor plays a role. UniProt data was accessed via the online database (https://www.uniprot.org/uniprotkb/?facets=reviewed%3Atrue&query=%2A), filtered for reviewed entries. The columns `Cofactor` and `EC Number` were used to identify which cofactors are associated with which ECs.

After compiling cofactors across each database, we created a master list of cofactors used in this study. Some cofactors were excluded or grouped together. The UniProt database in particular contained several named cofactors that we excluded or grouped with other cofactors (Table S13). For each cofactor, we created a regular expression (regex) to identify mention of that cofactor or one of its synonyms (Table S14). We ran the regexes against KEGG Enzyme entries' `comment` fields and Expasy `rdfs:comment` fields to flag ECs that might have cofactor dependencies. We generated a spreadsheet with all potential mappings of EC-cofactor dependencies (based on the aforementioned data from Expasy, PDBe, UniProt, and KEGG) and manually verified them (See Table S15).

We also ran our regexes against KEGG Compound names in order to identify cofactors that have corresponding KEGG Compound IDs. In some cases, cofactors did not have any corresponding KEGG Compound IDs, or several KEGG Compound IDs mapped to a single cofactor. For this reason, we assigned each cofactor a single "ZID" (analogous to a KEGG Compound ID) that we could use to modify KEGG Reaction definitions to account for implicit cofactor dependencies (Table S12).

#### Addition of missing reactions

##### Oxygen-producing reactions and phylloquinone-dependent photosynthesis

We removed all $O_2$-producing reactions except for the water oxidizing complex (R09503), which relies on plastoquinone as the electron acceptor. We found that the only way to produce plastoquinone (the photosystem II coenzymes in the water oxidizing complex) was via homogentisate, using 4-hydroxyphenylacetate 1-monooxygenase (R02514/R02516), 3-hydroxyphenylacetate 6-hydroxylase (R02515), or hydroxyphenylpyruvate dioxygenase (R02521), which all use dioxygen as a co-substrate. This suggests that homogentisate production necessarily requires dioxygen. However, deletion of *ppd* (*slr0090*) hydroxyphenylpyruvate dioxygenase from *Synechocystis* sp. PCC 680 proved to be non-essential for plastoquinone biosynthesis [1], suggesting there are additional uncharacterized biosynthetic routes for producing homogentisate. Rather than assume plastoquinone (and thus oxygenic photosynthesis) depends on dioxygen for cofactor biosynthesis, we assumed that primitive versions of oxygenic photosynthesis could use alternative quinones. Since the charged version of P680 ($P680^*$) has a low reducing potential, we reasoned that phylloquinone could serve as a potential alternative electron acceptor [2]. We thus added a reaction (photosyn_R09503_vX) that uses phylloquinone instead of plastoquinone as the electron acceptor. We also required the chlorophyll precursor magnesium protoporphyrin IX 13-methyl ester (C04536) to be present before enabling the reaction.

##### L-aspartate oxidase

In KEGG, iminoaspartate (C05840) can be synthesized from aspartate (C00049) using either $NAD(P)^+$ or $O_2$ as an oxidant by aspartate dehydrogenase (1.4.1.21) or L-aspartate oxidase (1.4.3.16), respectively. However, in anaerobic conditions, L-aspartate oxidase can use fumarate as an oxidant [3]. Thus, we added a reaction producing iminoaspartate from L-aspartate with fumarate as the electron acceptor.

##### Protein synthesis reaction

We constructed a reaction to synthesize the KEGG metabolite C00017 representing a generic protein substrate using all 20 common amino acids as co-substrates.

##### Ferredoxin synthesis reactions

We modeled the synthesis of ferredoxins (C00138, C00139, C22154, C22150, C22151) using an ancestral model of ferredoxin [4,5], where five amino acids (Gly, Asp, Ala, Ser, and Cys) were proposed to be the protein scaffold for primitive ferredoxins. Exclusion of cysteine in this amino acid set, or requiring all 20 common amino acids for ferredoxin function, did not alter scope composition or size.

**Acyl-carrier protein synthesis reaction**

We constructed a reaction to model the synthesis of acyl-carrier protein (C00229) using the protein metabolite (C00017) and 4'-Phosphopantetheine (C01134).

**Thioredoxin/Thiol Protein/Protein sulfide/Sulfur carrier synthesis reactions**

The reduced and oxidized thioredoxins (C00343/C00342), thiol protein (C15814), and sulfur carrier protein (C15810) were produced directly from the protein metabolite C00017 because cysteine is required for the production of the protein metabolite.

**Flavodoxin/flavoprotein synthesis reaction**

We synthesized reduced and oxidized flavodoxin (C02745,C02869) and flavoproteins (C04253,C04570) using the protein metabolite (C00017) and FAD (C00016/Z00017).

**Cobalamin, corrinoid, and Cobamide biosynthesis reactions**

To enable cobalamin production, we had to gap-fill the production of cobyrinate (C05773) directly from cobalt-precorrin 8 (C11545) with no co-substrates. To synthesize corrinoid (C06021), we required the production of a generic heme compound (Z00025), cobalt (Z00006), and protein (C00017). For cobamide cofactor variants used in Wood-Ljungdahl pathway there was a missing reaction for amending specific lower ligands. We thus added a pseudo reaction synthesizing the 5-methoxybenzimidazolylcobamide cofactor from 5-methoxybenzimidazole (C22450) and the corrinoid protein (C06021).

**Quinone synthesis reactions**

Generic quinones (C15602) were producible from either p-benzoquinone (C00472) or o-benzoquinone (C02351). Since we cannot ascribe structural details to the specific quinones used in reactions annotated as using C14602, we assumed that the simplest quinone species, like benzoquinone, are sufficient to enable reactivity.

**[E2 protein]-L-lysine and [GCSH]-L-lysine reaction(s)**

Lipoyl-carrier protein component [E2 protein]-L-lysine (C22158) and the glycine cleavage system component [GCSH]-L-lysine (C22157) are key substrates for lipoyl-dependent reactions. We modeled the synthesis of each from the protein metabolite (C00017), which requires lysine (C00047).

##### Polyprenyl diphosphate synthesis

Polyprenyl diphosphate (C05847) is a key intermediate in the biosynthesis of various quinones, but is a disconnected generic metabolite in KEGG. We modeled synthesis of this cofactor directly from a prenyl diphosphate precursor, dimethylallyl diphosphate (C00235), with no other co-substrates.

##### Ferricytochrome synthesis reactions

We modeled the synthesis of ferricytochrome metabolites (C00125, C00126, C18233, C18234, C00999, C00996, C00924, C01070, C01071, C18233, C18234) from the generic protein metabolite (C00017), the generic iron metabolite (Z00015), and the generic heme metabolite (Z00025).

##### H4MPT synthesis reaction

Tetrahydromethanopterin, or H4MPT (C05927) was proposed to be synthesized from N-[(7,8-Dihydropterin-6-yl)methyl]-4-(beta-D-ribofuranosyl)aniline 5'-phosphate (C20562).

##### Prenylated FMN synthesis reaction

Synthesis of  the cofactor prenylated FMN (Z00057) required reduced FMN (C01847) and prenyl phosphate precursors (C00235).

##### Azurin synthesis reaction

We modeled the synthesis of azurins (C05357,C05358) using the protein metabolite (C00017) and the generic copper metabolite (Z00070)

##### Methanophenazine synthesis reaction

Methanophenazine (C11903,C11904) is a key cofactor for methanogens with cytochromes and we modeled its synthesis using all-trans-pentaprenyl diphosphate (C04217) and phenazine (C21476).

##### PQQ synthesis reaction

Pyrrolo-quinoline quinone (PQQ; C00113) can be modeled as a radical SAM mechanism on a tyrosine residue [6]. We thus required S-adenosylmethionine (C00019), glutamate (C00025), and tyrosine (C00082) for synthesis of this cofactor.

##### Methanofuran synthesis reaction

We modeled the synthesis of methanofuran (C00862) as occurring directly from the precursor (4-{4-[2-(gamma-L-Glutamylamino)ethyl]phenoxymethyl}furan-2-yl)methanamine (C21070).

##### Polyphosphate synthesis reaction

We modeled the synthesis of the generic metabolite polyphosphate (C00404) from triphosphate (C00536).

#### Model of ancient succinate semialdehyde production

We found that the inclusion of the organic seed molecules listed in Table S4 were required for expansion to more than 4,000 compounds, and that including just pyruvate resulted in very small networks (scope size of 373 compounds). Including succinate semialdehyde in the seed set produced a network of *n*=4089 compounds, suggesting that production of succinate semialdehyde was limiting the expansion. We hypothesized that a reaction producing succinate semialdehyde from succinate and molecular hydrogen could enable expansion from pyruvate as the sole source of organic carbon. We thus added the following reaction to the network:

$$\text{Succinate} + H_2 \rightarrow \text{Succinate semialdehyde} + H_2O$$

This reaction has an estimated free energy at standard molar conditions $\Delta_r G'^{\circ} = 1.0 \pm 7.3$ [kJ/mol] at pH=7.0, pMg = 3.0 and an ionic strength of 0.25M in aqueous conditions [7], We found activities within the feasible ranges listed in the Methods section of the main text that make this reaction thermodynamically feasible. We next repeated the expansion with just pyruvate as the organic carbon source and produced a network of *n*=4093 compounds.

#### A note on potential ways to improve the model in future research

The model presented in the paper can be improved upon in future work. First, ~50% of the compounds in the KEGG database are still not reachable, suggesting that many reactions are not included in the KEGG database, which may be overcome by incorporating additional biochemical databases or a much broader scope of hypothetical reactions (Fig. S1). Second, a large fraction of biochemical reactions have unknown standard molar free energies, which may be addressable using novel quantum mechanics-based methods for free energy estimation. Finally, whereas our model explicitly considers dependencies on small organic and inorganic cofactors based on known enzyme mechanisms, future models can incorporate more complex rules encoding dependencies for specific metals or protein-based catalysts.

#### The geologic record and metabolic network expansion

Each extant carbon fixation pathway is accompanied by a characteristic separation of $^{13}C$ from $^{12}C$ in the initial inorganic substrates (e.g., $CO_{2\,(g)}$, $CO_{2\,(aq)}$, $HCO3^{-}_{(aq)}$) relative to the ultimate metabolic products that make up bulk biomass [8]. The carbon isotope compositions of carbonate minerals and organic matter in ancient rocks reflect these 'fractionations', albeit as affected by the isotopic consequences of carbon transformations in the water column, sediments, and rocks [9]. To a first order approximation, such carbon isotopic comparisons are exercises in identifying extreme values that are potentially inaccessible to carbon fixation pathways that are less isotopically selective (e.g., rTCA) but plausibly produced by others (e.g., the Calvin cycle, reductive acetyl-CoA pathway).

Carbon isotope compositions of graphite grains from rocks and mineral inclusions >3.7 to 4.1 billion years old [10] are enriched in $^{12}C$ relative to carbon in the bulk Earth, and have been interpreted as biogenic in origin. Assuming a solid-earth source of $CO_2$, these low $^{13}C$-$^{12}C$ ratios would seem to rule out operation of all extant carbon fixation pathways except for either the Calvin cycle or the reductive acetyl-CoA pathway (or both) on the earliest Earth [11]. While the carbon isotope differences between $CO_2$ and $CH_4$ trapped in fluid inclusions in ~3.48 billion-year-old hydrothermal cherts has been interpreted as the result of hydrogenotrophic and/or acetoclastic methanogenesis [12], they have also been interpreted as the high-temperature abiotic $CO_2$ reduction [13]. We stay agnostic to this debate and do not consider these measurements here. The first paired carbon isotope measurements from carbonate minerals and associated organic carbon come from slightly younger rocks (~3.43 billion years ago, [14]) and are also consistent with carbon fixation by the extant Calvin cycle and reductive acetyl-CoA pathway. In fact, locale-specific and global compilations of carbon isotope measurements indicate that this is the case until ~2.9 billion years ago, where the carbon isotopic difference between carbonate minerals and organic carbon starts to exceed the range produced by either the extant Calvin cycle or reductive acetyl-CoA pathway. This feature reaches its extreme around ~2.7 billion years ago in sediments from shallow water environments and ~2.6 billion years ago in deeper water sediments [15,16], and is attributed to newfound production of $^{12}C$-enriched methane and subsequent incorporation of methane-derived carbon into biomass through methanotrophy and/or aerobic methane oxidation [17,18]. While this has been proposed to reflect the ecological availability of oxidants leading up to the great oxidation event, an equally compelling possibility is that it reflects the near coincident origination of methanic metabolisms (Fig. 4d). If this is the case, then the carbon isotope record prior to ~1 billion years of Earth's history could largely be a result of the something like the extant Calvin cycle, with variants of the reductive acetyl-CoA pathway that are associated with methanogenesis [19] appearing only ~2.9 billion years ago. Such an ordination is consistent with the metabolic expansion proposed here (see Fig. 4c in main text). Importantly, in contrast with previous models [19], the model presented of metabolic evolution in the main text predicts an early emergence of methanol-driven, prior to hydrogenotrophic methanogenesis. Future efforts to resolve this timing could be explored through physiologically controlled experiments on carbon isotope fractionation among variants of methanogenesis pathways.

### Extended Data Figures

#### Extended Data Fig. 1

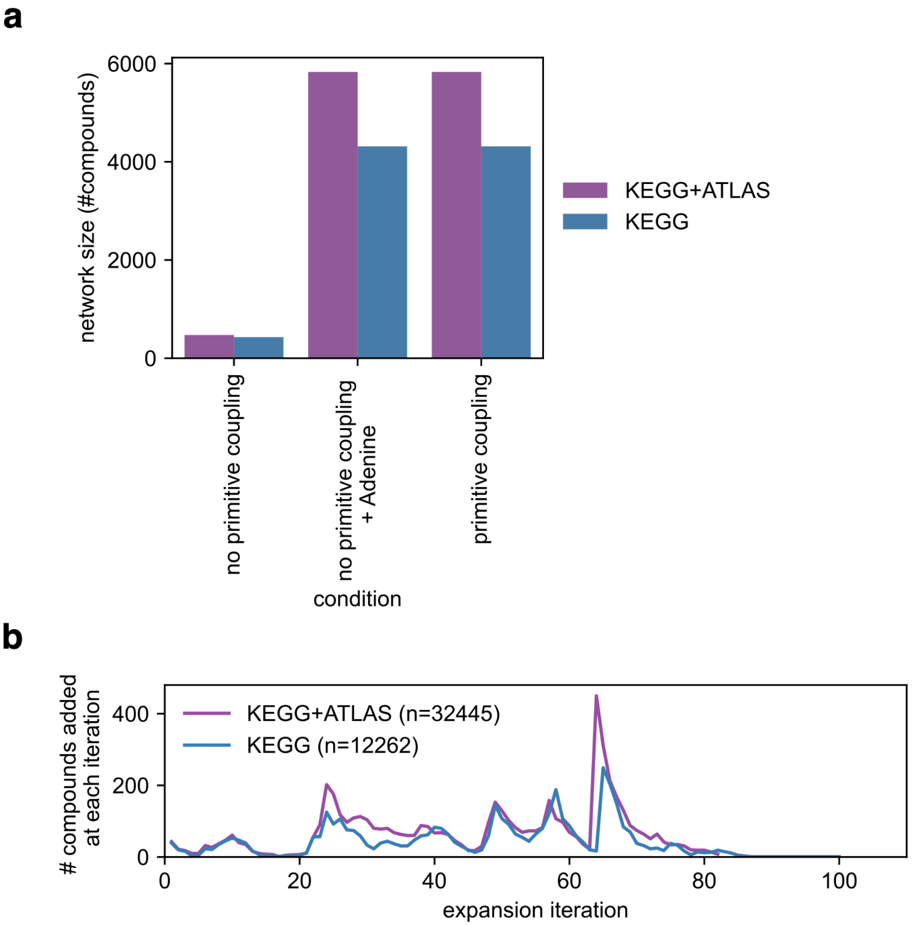


**Extended Data Fig. 1: Network size and trajectory of metabolic expansion with hypothetical biochemistry.** We performed network expansion using the same seed set as shown in Fig. 1 in the main text, but added 20,183 additional reactions from the ATLAS database [20,21]. (**a**) The network sizes (number of compounds produced) for the expansion using KEGG only (blue) or KEGG plus the ATLAS database (purple) with either (i) no primitive coupling model, (ii) no primitive coupling, but added adenine to the seed set, or (iii) added primitive phosphate coupling. **(b)** The number of compounds produced at each iteration (*y*-axis) for the network composed of both KEGG and ATLAS reactions with primitive coupling (purple line) results in a punctuated structure similar to that of the KEGG reaction network with primitive coupling (blue line; see also Fig. 2b).

**Extended Data Fig. 2**

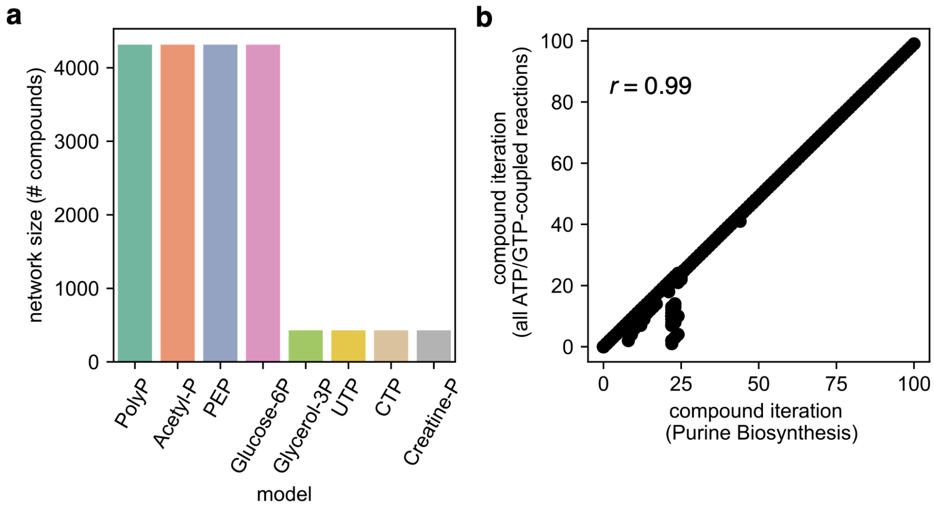


**Extended Data Fig. 2: Scope sizes with alternative models of phosphoryl donor promiscuity: (a)** We constructed hypothetical *de novo* purine biosynthesis pathways with one of 6 potential phosphoryl donors in KEGG ($PP_i$ = pyrophosphate, Acetyl-P= acetyl phosphate, PEP=phosphoenolpyruvate, Glucose-6P= glucose-6-phosphate, Glycerol-3P=glycerol-3-phosphate, UTP=uridine triphosphate, CTP=cytidine triphosphate, and Creatine-P=creatine phosphate). The ensuing network sizes are plotted on the *y*-axis. Note that pyrimidine triphosphates did not enable expansion above 1000 metabolites. (**b)** A comparison between including primitive pyrophosphate-coupled purine synthesis versus substituting all ATP/GTP-coupled reactions with pyrophosphate alternatives (*n*=523). A scatterplot for all compounds emerging in both expansions, where the *x*-axis denotes the iteration number for which a compound appears in the primitive purine biosynthesis model, and the *y*-axis corresponds to the iteration when the same compound emerges in the model where all ATP/GTP-coupled reactions have pyrophosphate alternatives. The positive correlation coefficient (Pearson's $r$=0.99, $P<10^{-10}$) denotes a strong similarity in expansion trajectories for both models.

**Extended Data Fig. 3**

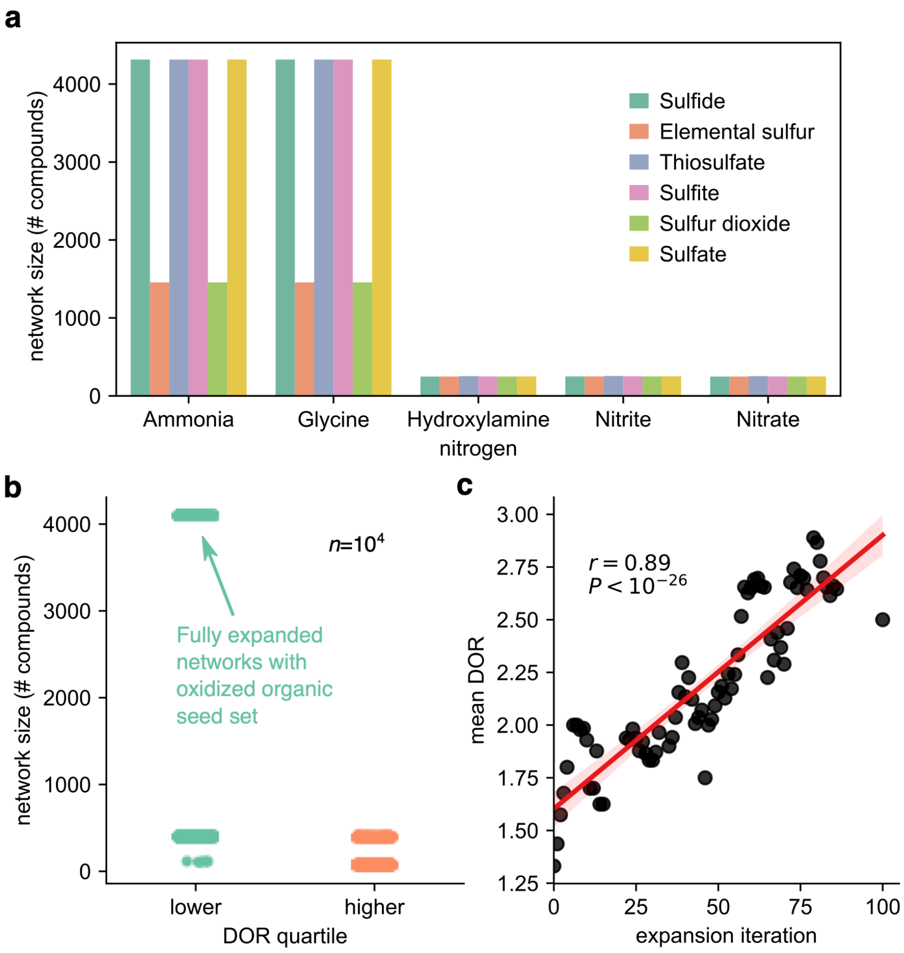


**Extended Data Fig. 3: Expansion with alternative nitrogen, sulfur, and carbon sources at various redox states changes scope and trajectory of metabolic evolution. (a)** We altered the nitrogen ($x$-axis) and sulfur sources (color) of the seed set (listed Table S4) and re-ran the expansion. The size of the network (i.e., number of compounds in the scope) is plotted on the $y$-axis. Only expansions with fully reduced nitrogen led to large networks, while several sulfur sources (sulfate, sulfite, thiosulfate, and sulfide) could lead to large networks. (**b**) We varied the composition of the organic molecules by randomly sampling sets of $N$=21 compounds from a ranked distribution based on computed the degree of reduction (DOR). As described before [22,23], the DOR for molecule $C$ is calculated using the following formula: $y/x$; $x\mathrm{CO_2} + y\mathrm{H_2} = C + z\mathrm{H_2O}$. We either sampled compounds in the bottom quartile of the distribution (lower, green dots) or from the top quartile (higher, orange dots). We found that expansion to >4000 compounds required seed compounds that had a low DOR, which consists of more oxidized carbon sources. **(c)** The average DOR for molecules produced at each iteration ($y$-axis) was plotted against the expansion iteration ($x$-axis); these variables exhibit a significant correlation (Pearson's $r = 0.89$, $P<10^{-26}$).

**Extended Data Fig. 4**

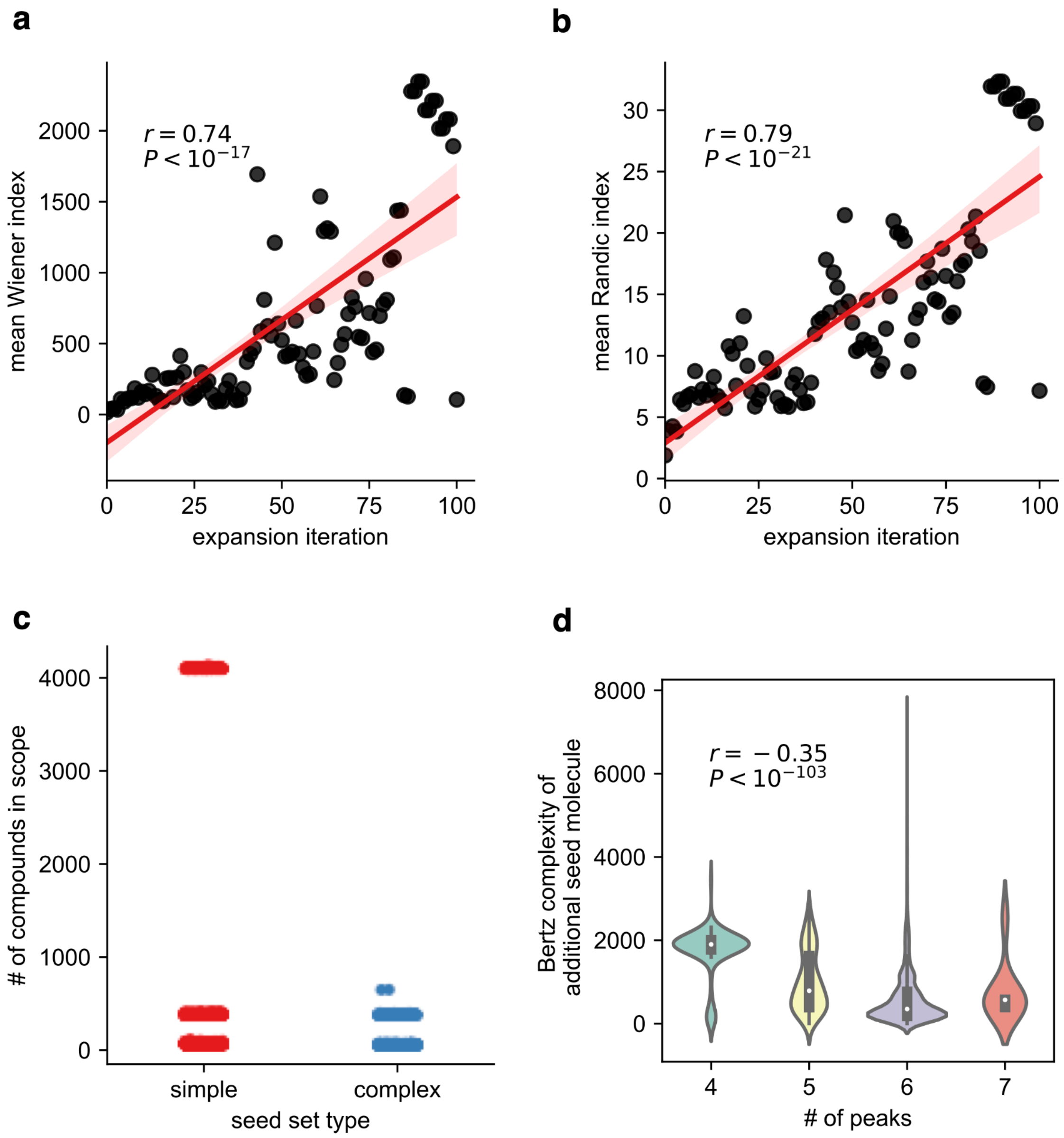


**Extended Data Figure 4. Molecular complexity and metabolic network expansion. (a)** The average Wiener topological index [24] (*y*-axis) at each expansion iteration (*x*-axis). **(b)** Same as **(a)** but for the Randic topological index [25]. (c) We sampled 21 organic molecules either at the top (blue dots, "complex") or bottom (red dots, "simple") quartile of molecules in terms of Bertz molecular complexity and re-ran network expansion. Only expansion with low complexity seed sets (red dots) led to expansions >4000 compounds. (**d**) For every compound we could compute a Bertz complexity metric for ($n$=3543), we added that compound into the original seed set and re-ran network expansion. We next quantified the number of peaks in the expansion trajectory using the `find_peaks` function in the `scipy.signal` module with parameters height=0, distance=5, and prominence=30. We found that the number of peaks was inversely correlated with the Bertz complexity of the additional seed molecule (Pearson $r$=-0.35 with $P < 10^{-103}$).

**Extended Data Fig. 5**

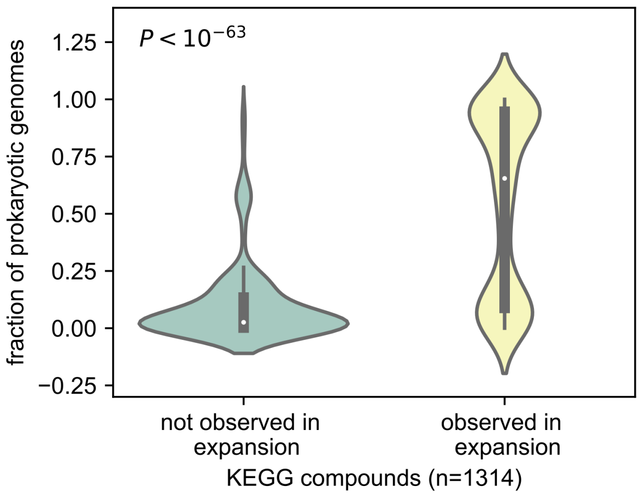


**Extended Data Fig. 5. Microbial genomes encode pathways that use compounds in expansion scope at higher rates than compounds outside of expansion scope.** We took 1,312 KEGG compounds that are intermediates in 377 KEGG modules, computed the proportion of prokaryotic genomes that use each compound as an intermediate in at least one KEGG module ($y$-axis), and grouped each compound based on whether the compound was observed in the expansion scope or not ($x$-axis). We plotted the distributions as violin plots and found that compounds observed in the expansion scope ($n$=1,024) were much more prevalent in microbial genomes than compounds not observed in the expansion scope ($n$=288) (Mann-Whitney $U$ test: $P < 10^{-63}$)

**Extended Data Fig. 6**

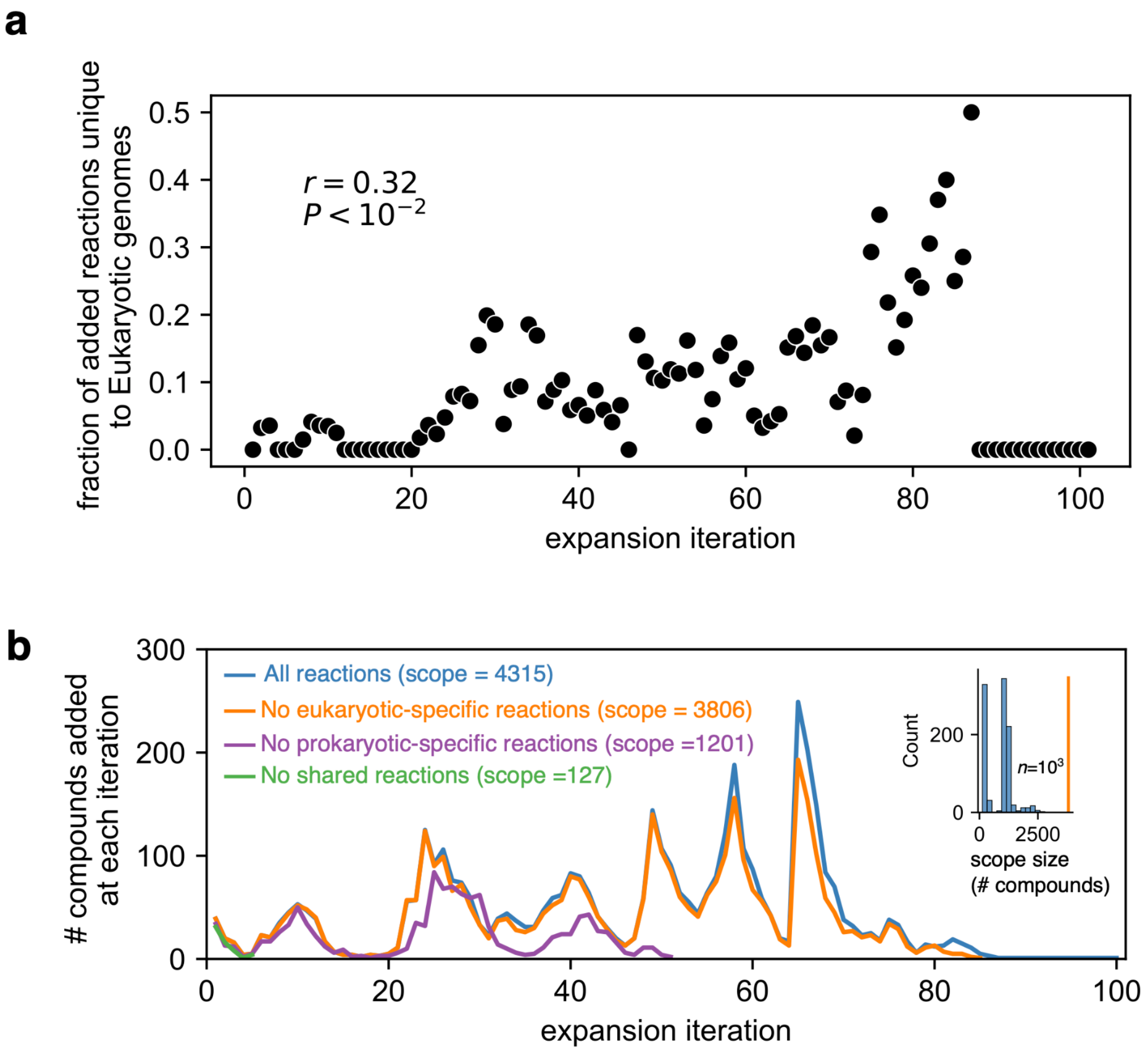


**Extended Data Figure 6. Utilization and dependence of eukaryote-specific reactions during expansion.** (a) We identified 1638 reactions that were uniquely mapped to eukaryotic genomes, and plotted the proportion of reactions added during each expansion iteration that were eukaryote-specific (*y*-axis). We find a weak positive correlation between the relative proportion of eukaryote-specific reactions with expansion iteration (Pearson's $r$=0.32, $P$<$10^{-2}$). (b) We removed these reactions, performed the expansion, and found that the expansion scope consisted of 3806 compounds (88% of the reported expansion scope in the main text), with similar punctate structure. We found that removal of these 1638 reactions led to a particularly large network compared to expansions after randomly removed reaction sets: we sampled $10^3$ sets of reactions (all of size 1638) to remove and re-ran network expansion. For all randomly chosen sets, all networks were smaller than the eukaryotic depleted network (inset; the orange line denotes the scope size of the network after removing Eukaryote-specific reactions).

**Extended Data Fig. 7**

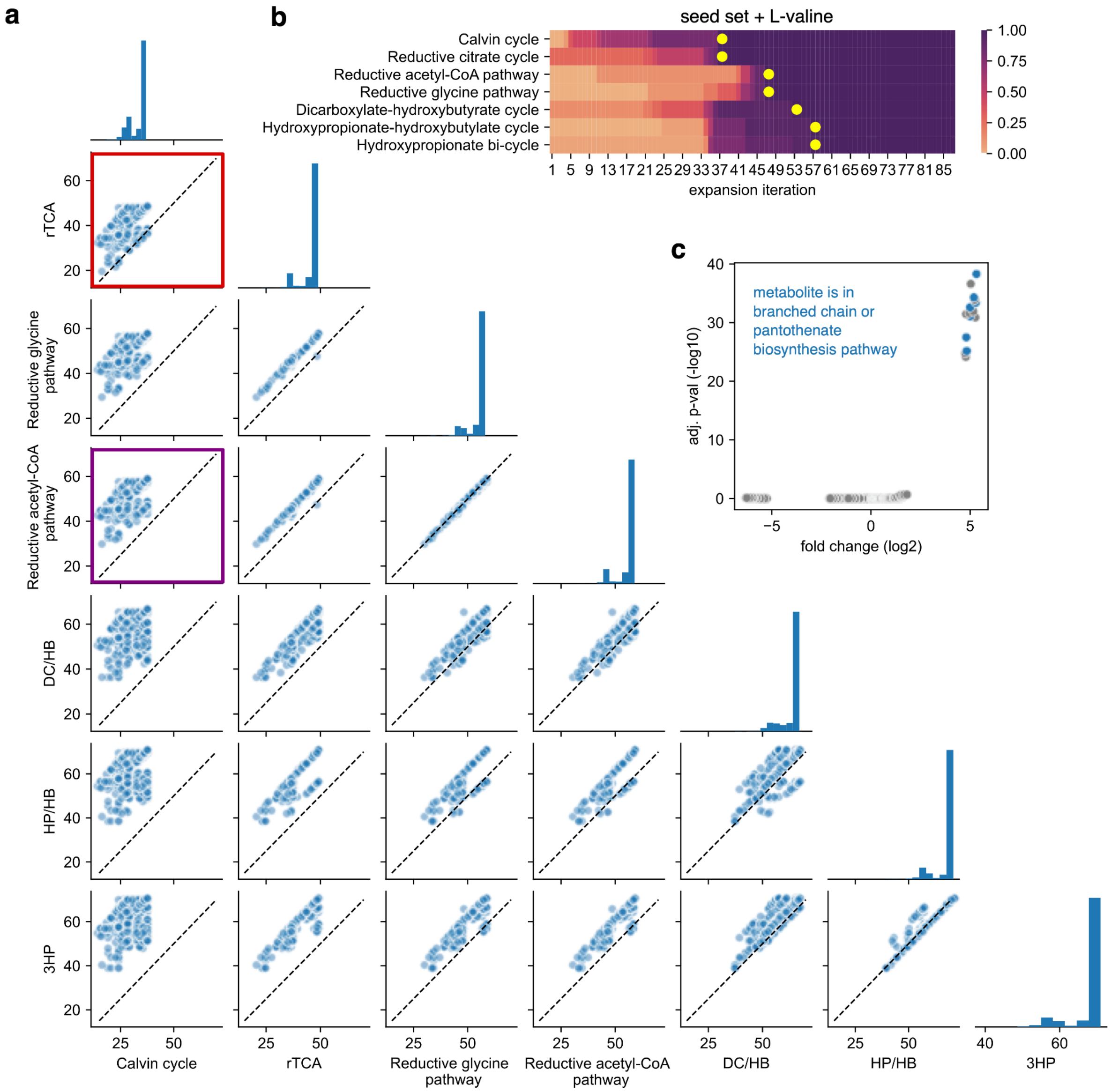


**Extended Data Fig. 7. Sensitivity of carbon fixation pathway timing to seed set variations.** **(a)** We randomly sampled 10 additional molecules from the set of KEGG compounds with less than 6 carbons (*m*=1805), repeated the expansion, computed the iterations when each pathway became feasible (see Fig 4c,d in the main text), and repeated the process $10^4$ times. (**a**) A pairwise scatterplot between the iteration of completion for each carbon fixation pathway, where the dashed line is the line of unity. The red box highlights the comparison between the Calvin cycle and the reductive citric acid (rTCA) cycle, showing that a subset of conditions led to the Calvin cycle and rTCA emerging at the same iteration. In contrast, the purple box highlights the comparison between the Calvin cycle and the reductive acetyl-CoA pathway, showing that for all expansions, the Calvin cycle proceeds the extant reductive acetyl-CoA pathway during the expansion. (**b**) A heatmap showing the pathway completion for an expansion with the original seed set, plus valine, which results in both the Calvin cycle and rTCA cycle emerging at the same iteration due to the accelerated production of Coenzyme A (CoA). (**c**) A volcano plot for each molecule randomly added as a seed compound. For each metabolite, we computed the enrichment of that metabolite in the seed set for expansions that led to identical completion iterations between the Calvin and rTCA cycle (*x*-axis), and computed the statistical significance of this enrichment (*y*-axis) using a Fisher's exact test. Metabolites that are over-represented in expansions where the Calvin cycle emerged at the same iteration as the rTCA cycle fall within the top right corner of the plot. Several of these metabolites are involved in either branched chain amino acid or pantothenate biosynthesis, which are both key precursors for Coenzyme A.
